# The *Drosophila* cardiogenic transcription factors Myocyte enhancer factor-2 and Tinman contribute to heart lumen enlargement through direct activation of the collagen gene *Multiplexin*

**DOI:** 10.1101/2023.06.28.546941

**Authors:** TyAnna L. Lovato, Danielle Ryan, Cayleen Bileckyj, Christopher A. Johnston, Richard M. Cripps

## Abstract

During embryogenesis, the Drosophila heart forms a lumen, the posterior region of which is increased in diameter and corresponds to the heart proper. To identify the transcriptional control of this morphogenetic process, we analyzed the formation and enlargement of the heart lumen in mutants for the myogenic transcription factor gene *Myocyte enhancer factor-2* (*Mef2*). We found that *Mef2* contributes to both lumen formation and lumen expansion, the latter through a requirement for both *Mef2* and the cardiogenic gene *tinman* (*tin*) to activate the collagen gene *Multiplexin* (*Mp*). To determine if Tin and MEF2 act directly upon the *Mp* gene, we identified an enhancer whose activity recapitulates the cardiac expression of *Mp*. This enhancer contains binding sites for both Tin and MEF2 and is activated in tissue culture by MEF2 but not Tin. We did not observe synergistic activation of the enhancer when the factors were in combination, despite documenting a direct physical interaction between Tin and MEF2 in vitro. In vivo, the Tin sites are required for normal enhancer activity, whereas mutation of the MEF2 sites results in expanded expression of an enhancer-*lacZ* reporter, suggesting that transcriptional repression may also contribute to regulation of *Mp*. Our studies underline how transcription factors must utilize combinatorial interactions to achieve organ-specific and region-specific patterns of gene expression and cell morphogenesis.

## Introduction

Early heart development in animals arises from the convergence of bilateral groups of cardial cells that interact to form a hollow contractile tube through which hemolymph or blood is pumped (see Souidi & Jagla, 2021 for review). Research in Drosophila has determined that cardiac tube formation arises from a carefully orchestrated series of events involving cardiac signals, receptors, and extracellular matrix. Early studies demonstrated that cardiac cell migration required the secreted glycoprotein Slit and its receptor Robo (Santiago-Martinez et al 2006), and these factors are subsequently required to regulate cell adhesion during lumen formation (Qian et al 2005; Santiago-Martinez et al 2006; Medioni et al 2008). Slit/Robo accumulation and function are in turn regulated in a complex manner: the genes *held out wings* and *Dystroglycan* are necessary for normal Slit localization (Medioni et al 2008); and the localizations of Slit, and in some instances Robo, are regulated by additional mechanisms including Integrin accumulation (Vanderploeg and Jacobs, 2012). Moreover, the collagen Multiplexin (Mp) potentiates Slit/Robo signaling to enhance lumen size in the posterior-most two segments of the embryonic heart (Harpaz et al 2013). Additional factors also are required for lumen formation, including the transmembrane proteoglycan Syndecan (Knox et al 2011), the transmembrane receptor Unc-5 (Albrecht et al 2014), and the adaptor protein Talin (Vanderploeg and Jacobs 2015).

There is compelling evidence that mechanisms of cardiac morphogenesis are conserved in diverse animal species, and in several cases this includes lumen formation. In zebrafish, members of both the Slit and Robo gene families are expressed during lumen and chamber formation, and morphants for *Slit2, Robo1* or *Robo4* have severe defects in cardiac cell migration and lumen formation (Fish et al 2011). Similarly in mice, knocking out Slit or Robo orthologs results in abnormalities including ventral septal defects, aberrant cardiac valve formation, pericardial abnormalities, and vein defects (Mommersteeg et al 2015; Zhao et al 2022).

More importantly, recent studies have shown that derangement of these cardiac signaling processes may underlie some forms of human congenital heart disease. Mutations in *Robo1* are associated with congenital heart defects including atrial septal defects and Tetralogy of Fallot (Kruszka et al 2017); and mutations in *Robo4* are associated with aortic valve defects and aortic aneurysm (Gould et al 2019).

These studies underline the progress that has been made in identifying the structural genes controlling heart lumen formation, yet there is still much to learn of how this process is regulated on a more global scale. Specifically, what transcription factors contribute to expression of these structural genes in the heart? Using a genetic interaction approach, Qian et al (2011) demonstrated that the Rho GTPase cdc42 functions with the cardiogenic factor Tinman (Tin) to regulate normal heart growth and function, and Vogler et al (2014) showed that cdc42 regulates heart cell polarity and lumen formation. In a related work, Asadzadeh et al (2015) showed that u*nc-5* is a direct transcriptional target of Tin in the heart, identifying for the first time a direct connection between a cardiac transcription factor and a regulator of lumen formation.

Here, we determine the role of the myogenic regulatory factor Myocyte enhancer factor-2 (MEF2) in heart lumen formation and enlargement in Drosophila. We demonstrate that lumen formation is attenuated in *Mef2* null embryos, and these mutants display a failure of lumen enlargement in the posterior heart region. We show that the latter phenotype arises at least in part through a requirement for both MEF2 and Tin in *Mp* transcription, and we define and characterize a *Mp* enhancer that binds directly to MEF2 and Tin in vitro, and whose activity is dependent upon the integrity of these sites. Our studies begin to uncover the transcriptional mechanisms that orchestrate heart lumen formation and provide insight into how cardiac defects might arise from transcription factor dysfunction.

## Materials and methods

### Drosophila methods

Drosophila were maintained at 25°C for all experiments. Gene names and symbols are as described at FlyBase.org. The *tinABD; TM3 lacZ/tin*^*346*^ line was from Dr. Rolf Bodmer (Sanford-Burnham-Prebys Institute). Transgenic lines were created by injecting plasmid DNAs (described below) into the Phi-C31 integrase line M{3xP3-RFP.attP}ZH-86Fb (Bischof et al 2007) (BDSC 24749). Some constructs were injected in-house, while other were injected by Rainbow Transgenic Flies, Inc. Transgenic lines were identified in the G1 generation after intercrossing the G0 adults, and homozygous lines were generated in the subsequent generation based upon stronger rescue of the white eye colour.

### DNA methods

Wild-type enhancer fragments from *Mp* (with the exception of Mp3x, Mp3y and Mp3z) were generated by PCR using sequence-specific primers that also contained attB sites. The attB-flanked PCR products were directly recombined into the *lacZ* reporter construct pDONR-lacZ-attB (Bryantsev et al 2012) using BP Clonase (ThermoFisher). Recombinant plasmids were sequenced to confirm that the inserted DNA was correct, and then prepared for injections through Qiagen midipreps. The constructs Mp3x, Mp3y, and Mp3z, plus mutant versions of Mp3z were created as gene synthesis fragments containing flanking attB sites (Integrated DNA Technologies Inc.) and then directly recombined into pDONR-lacZ-attB.

### Generation of anti-Tin antiserum

Tin antiserum was generated essentially as described by Vishal et al (2020). Briefly, the coding sequence of *tin* was cloned into the pEXP1-DEST vector (ThermoFisher V96001) utilizing Gateway Technology via pDONR221 (ThermoFisher 12536017), and purified as denatured protein. Two New Zealand White rabbits were injected at ten sites with 500 µl of Freund’s Complete Adjuvant (Sigma-Aldrich F5881) containing 500 µg of Tin protein. After four weeks, a booster injection of 100 µg of Tin protein in 200 µl of Freund’s Incomplete Adjuvant (Sigma-Aldrich 344291) was injected at four sites. After a further two weeks 50 ml of blood was drawn to check titer, and a second booster was administered a week later. Four weeks later the final blood draw was obtained. Serum was collected by allowing the blood to coagulate at room temperature for 30 minutes and then centrifuged at 2000 x g. The supernatant was transferred to a clean tube and analyzed by immunohistochemistry at a concentration of 1:1000. 50 ml of blood was collected for each final blood draw. The rabbits were not exsanguinated and were both adopted according to approved protocol. The protocol was IACUC approved (protocol number 18-200741-MC) prior to commencing the work.

### Embryo stains and analysis

Embryo stains were performed essentially as described by Patel (1994), with localized primary antibodies being detected either immunohistochemically using the ABC detection method (Vector Laboratories), or via immunofluorescence using Alexa-conjugated secondary antibodies (Thermo Fisher). Primary antibodies, sources and concentrations were: mouse anti-ß-Galactosidase, Promega Corp., 1:500; guinea pig anti-H15, James Skeath (Washington University at St Louis), 1:1000; rabbit anti-Tin, 1:1000 after pre-absorption; mouse anti-ßPS integrin, Developmental Studies Hybridoma Bank, 1:50; rabbit anti-Dystroglycan, Dr. Roger Jacobs (McMaster University), 1:250; and anti-MEF2, this laboratory (Vishal et al 2020), 1:1000. Secondary antibodies were either biotinylated goat anti-mouse (1:500) for immunohistochemistry (Vector), or Alexa-conjugated antibodies (1:2000) for immunofluorescence (Fisher Scientific).

In situ hybridization to embryos was carried out essentially as described by O’Neill and Bier (1994). An antisense *Mp* probe was synthesized from an *Mp* cDNA, using the Digoxigenin labeling kit (Sigma). Hybridized Digoxigenin-labeled probe was visualized with peroxidase-linked anti-Digoxigenin followed by NBT/BCIP substrate.

Stained embryos were observed and imaged using an Olympus BX61 compound microscope for immunohistochemically-stained samples, or using and Olympus Fluoview 3000 confocal microscope for fluorescently stained samples. At least ten embryos of each genotype were analyzed, and representative images are shown.

### Tissue culture experiments

Drosophila S2 cell co-transfections and quantitative ß-Galactosidase assays were essentially as described by Kelly Tanaka et al (2008). Expression plasmids were pPAc-Tin and pPAc-Mef2, with pPAc-pl (Drosophila Genomics Resource Center) as the empty vector control. The reporter construct was the same *Mp3D-lacZ* reporter used for generating transgenic animals. Data represent average of at least three transfections for each different expression plasmid.

### Protein purification and protein interaction studies

The MEF2 MADS plus MEF2 domain (codons 1-90) was cloned into pBH, containing an N-terminal TEV-cleavable hexahistidine tag, using 5’-BamHI and 3’-XhoI restriction sites. The Tin homeodomain (codons 301-360) was cloned into pMAL, containing an N-terminal MBP tag, using 5’-NdeI and 3’-XhoI restrictions sites.

All proteins were expressed in BL21(DE3) *E. coli* under induction of isopropyl β-d-1-thiogalactopyranoside (IPTG) and grown in standard Luria–Bertani broth supplemented with 100μg/ml ampicillin. Transformed cells were grown at 37°C to an OD_600_ ∼0.6 and induced with 0.2 mMIPTG overnight at 18°C. Cells were harvested by centrifugation (5000×*g* for 10 min), and bacterial pellets were resuspended in lysis buffer and flash-frozen in liquid nitrogen. Cells were lysed using a Branson digital sonifier and clarified by centrifugation (12,000×*g* for 30 min).

For hexahistidine-tagged MEF2, cells were lysed in N1 buffer (50mM Tris pH8, 500mM NaCl, 10mM imidazole) and coupled to Ni-NTA resin for 3 h at 4°C. Following extensive washing, proteins were eluted with N2 buffer (50mM Tris pH8, 500mM NaCl, 300mM imidazole). The hexahistidine tag was removed using TEV protease during overnight dialysis into N1 buffer. Cleaved products were reverse affinity purified by a second incubation with Ni-NTA resin and collection of the unbound fraction. Final purification was carried out using an S200-sephadex size exclusion column equilibrated in storage buffer (20mM Tris pH8, 200mM NaCl, 2mM DTT).

For MBP-tagged Tin, cells were lysed in lysis buffer (50mM Tris pH 8, 500mM NaCl, 2mM DTT) and coupled to amylose resin for 3 h at 4°C. Following extensive washing, proteins were eluted with elution buffer (50mM Tris pH8, 300mM NaCl, 2mM DTT, 50mM maltose). Final purification was carried out using an S200-sephadex size exclusion column equilibrated in storage buffer (20mM Tris pH8, 200mM NaCl, 2mM DTT).

Equivalent amounts of MBP or MBP-fused Tin bait constructs were absorbed to amylose agarose resin for 30 min at 4°C and washed three times to remove unbound protein. Subsequently, soluble untagged prey MEF2 protein was added for 1 h at 4°C with constant rocking in wash buffer (20mM Tris, pH 8, 120mM NaCl, 1mM DTT, and 0.2% Triton-X100). Reactions were then washed four times in wash buffer, and resolved samples were analyzed by Coomassie blue staining of SDS-PAGE gels. Gel shown in figure is representative of 3 independent experiments.

### DNA binding assays

Electrophoretic mobility shift assays were carried out using standard approaches, and reaction buffer as described by Gossett et al (1989). For the Tin shifts, purified native Tin-homeodomain (as described above) was used. For MEF2, protein was synthesized from pCDNA-Mef2 (Cripps et al 2004) in a coupled transcription-translation reaction using T7 RNA polymerase. Double-stranded DNA probes were generated by annealing single-stranded complementary oligonucleotides, that each contained a 5’-GG extension; this extension could be radioactively labeled by filling in the overhangs using 32P-dCTP and Klenow enzyme (NEB). Sequences of the +-strand oligonucleotides used are described in Table 1.

**Table 1.**
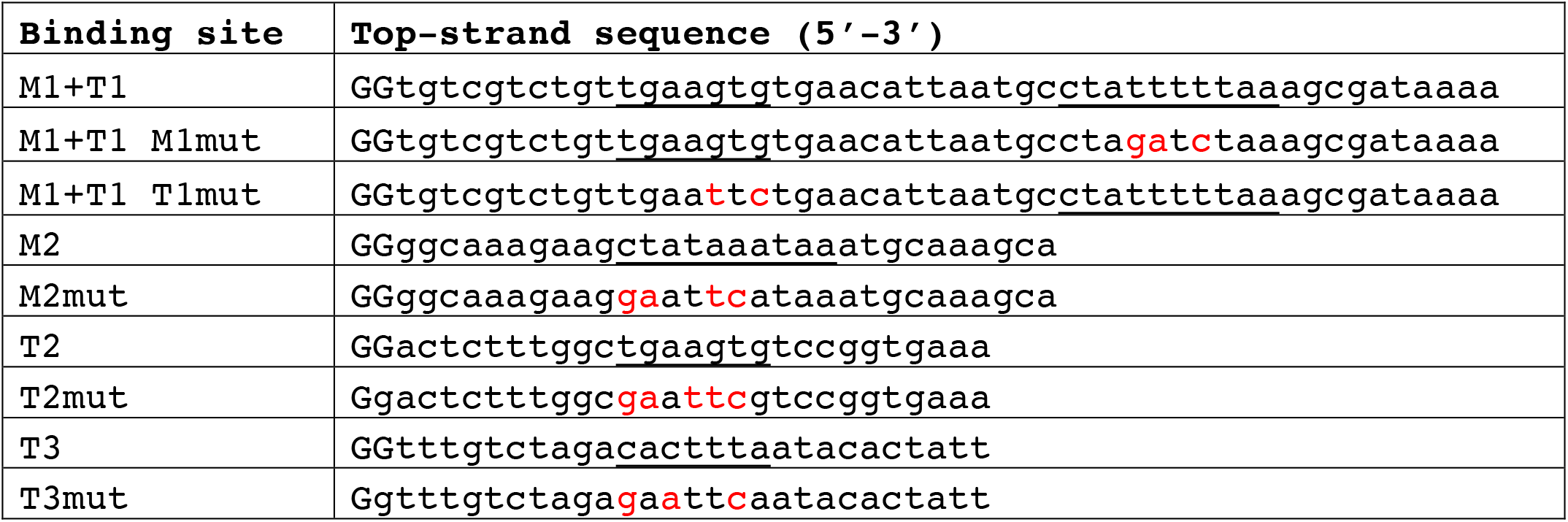
Top-strand sequences of probes used for DNA binding assays. The 5‘-GG dinucleotides are added for radioactive labeling (see text); consensus binding sequences are underlined; mutated sequences are shown in red.

## Results

### MEF2 is required for heart lumen formation and expansion of the heart proper

To gain insight into the transcriptional control of heart lumen formation and expansion, we characterized the heart in *Mef2* mutant embryos at stage 16 of development. In wild-type embryos (Figure 1A, C), the bilateral cardial cells accumulated high levels of ßPS-Integrin on their luminal sides (Figure 1A-A’’’), and clear spacing between the stained cells was apparent in the posterior heart region (Figure 1C-C’’’, asterisk) indicating expansion of the lumen. By contrast, *Mef2* null embryos showed much weaker Integrin accumulation (Figure 1B-B’’’) and there was no evidence of lumen expansion (Figure 1D-D’’’). In addition, staining for Dystroglycan, whose accumulation also highlights the enlarged lumen, did not reveal normal morphogenesis of the heart (compare Figure 1C’’ with 1D’’). Given that Integrin is necessary for proper lumen formation (Vanderploeg and Jacobs, 2012), we conclude that *Mef2* mutant embryos do not form a proper lumen, and the heart also fails to increase in diameter in the most posterior segments.

**Figure 1:**
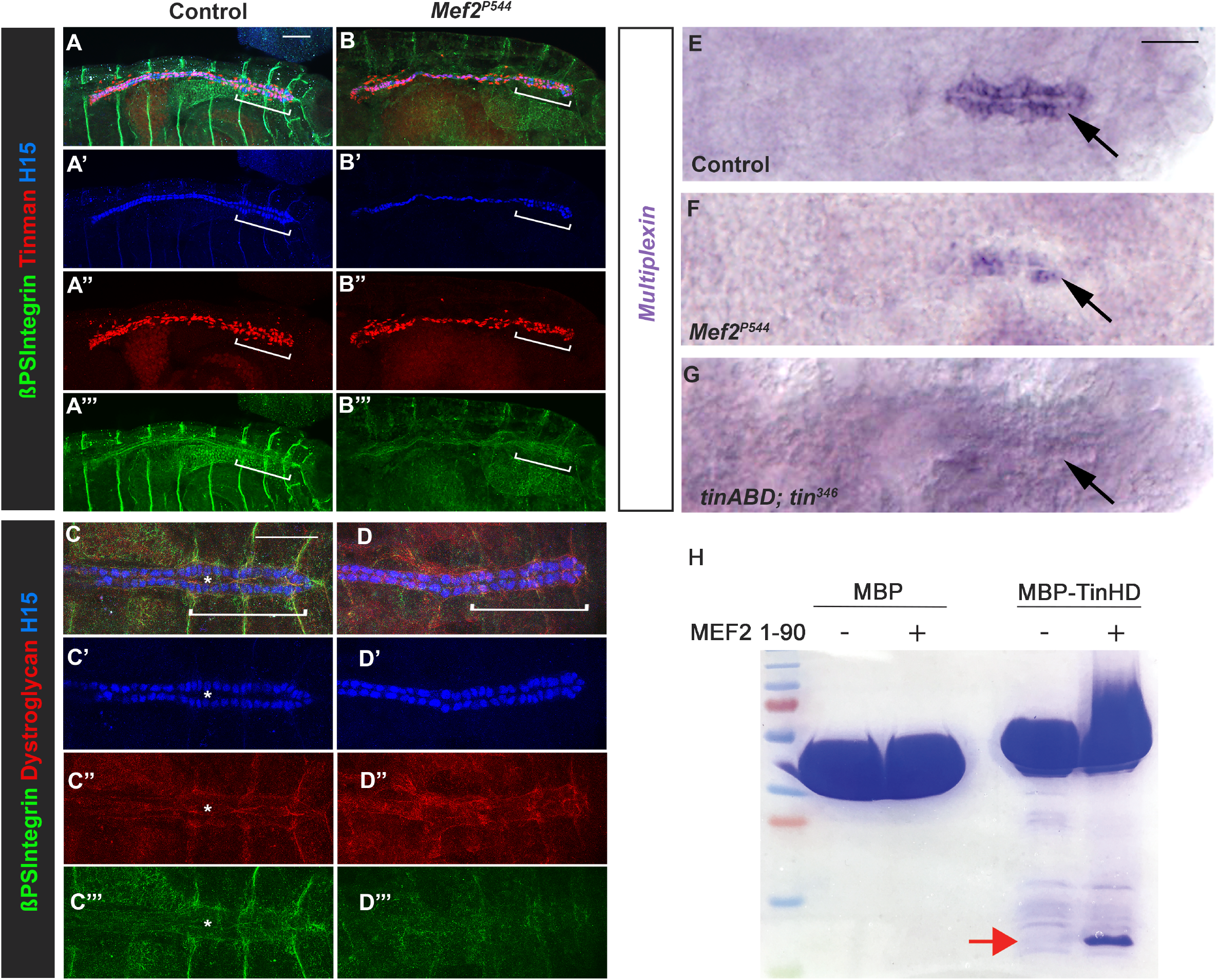
Roles for MEF2 and Tin in cardiac morphogenesis. **A**,**B:** The wild-type cardiac tube is broader in the posterior two segments corresponding to the heart proper (bracketed, **A-A’’’)**, but in *Mef2* null mutants the ordered rows of cardial cells observed in controls are irregular, and there is no obvious broadening of the heart in the posterior (**B-B’’’**). Integrin levels are strongly reduced in *Mef2* mutants compared to control (**A’’’, B’’’**). **C**,**D:** At higher magnification, the lumen of the heart can be observed in control animals (bracketed and asterisks in **C-C’’’**), but an enlarged lumen is never observed in *Mef2* mutants. **E-G:** Multiplexin expression is strongly detected in the posterior heart of control animals (**E**), but is reduced or absent in *Mef2* mutants (**F**) or late-*tin* mutants (**G**), respectively. Scale bars, 100µm for A-D, 50µm for E-G. Scale bars, 100µm for A-D, 50µm for E-G. **H:** MBP-TinHD, but not MBP alone, binds MEF2 (1-90) (red arrow).

### MEF2 and Tinman are required for Multiplexin expression in the heart proper

In mutants that lack late *tin* function, this enlargement of the heart tube was also not apparent (Zaffran et al 2006), suggesting that both *tin* and *Mef2* contribute to expansion of the cardiac lumen. In addition, Harpaz et al (2013) demonstrated that lumen enlargement was dependent upon expression of the collagen gene, *Multiplexin* (*Mp*), that shows highest cardiac expression in the myocardial cells of the posterior heart (Figure 1E). We then determined whether *Mp* expression was dependent upon either *Mef2* or *tin* function by staining control and mutant embryos for *Mp* transcripts. Here, we found that *Mef2* null mutant embryos always showed a reduction in *Mp* expression, although transcripts were still detected at low levels (Figure 1F). In late *tin* mutant embryos, we did not detect *Mp* transcripts (Figure 1G). These studies indicate that whereas *Mef2* is necessary for maximal levels of *Mp* expression, *tin* function is essential for *Mp* expression. Importantly, the loss of *Mp* transcripts in either of these mutant backgrounds can account at least in part for the failure of heart lumen expansion in the mutants.

In parallel to these experiments, we sought to determine if there is a physical interaction between the Tin and MEF2 transcription factors. This possibility was based both upon the requirement for each factor for full *Mp* expression, as well as the observed physical interaction between Nkx2.5 and MEF2C, the mammalian orthologs of Tin and MEF2, respectively (Vincentz et al 2008). We purified the MADS plus MEF2 domains of MEF2 (amino acids 1-90) using a cleavable hexahistidine tag and the homeodomain of Tin (amino acids 301-360) as an MBP fusion protein. MBP alone or MBP-Tin were then immobilized on amylose resin and subsequently incubated in the absence or presence of soluble MEF2. Following extensive washing, only the MBP-Tin reaction retained a detectable quantity of MEF2 protein that was visualized following elution and analysis by SDS-PAGE (Figure 1H).

Taken together, these studies showed that both Tin and MEF2 are required for *Mp* expression, and raised the possibility that they might collaborate in directly activating *Mp* transcription.

### Identification of an Mp cardiac enhancer

To test the hypothesis that both MEF2 and Tin directly regulate *Mp* expression, we sought to identify the *Mp* cardiac enhancer. The *Mp* gene spans ∼55kb of genomic DNA, has two distinct transcription start sites, and consists of numerous exons and large introns (Figure 2A); therefore, it was not immediately apparent where the cardiac enhancer might be located. Moreover, high-throughput studies have not revealed prominent interaction of either MEF2 or Tin with genomic DNA surrounding *Mp* (Sandmann et al 2007). This latter result may be due either to the relatively small size of the embryonic heart compared to the remainder of the embryonic mesoderm rendering the ChIP signal relatively weak, or to the existing ChIP data not covering the time of *Mp* activation in the heart, or to MEF2 and Tin not being direct regulators of *Mp*

**Figure 2:**
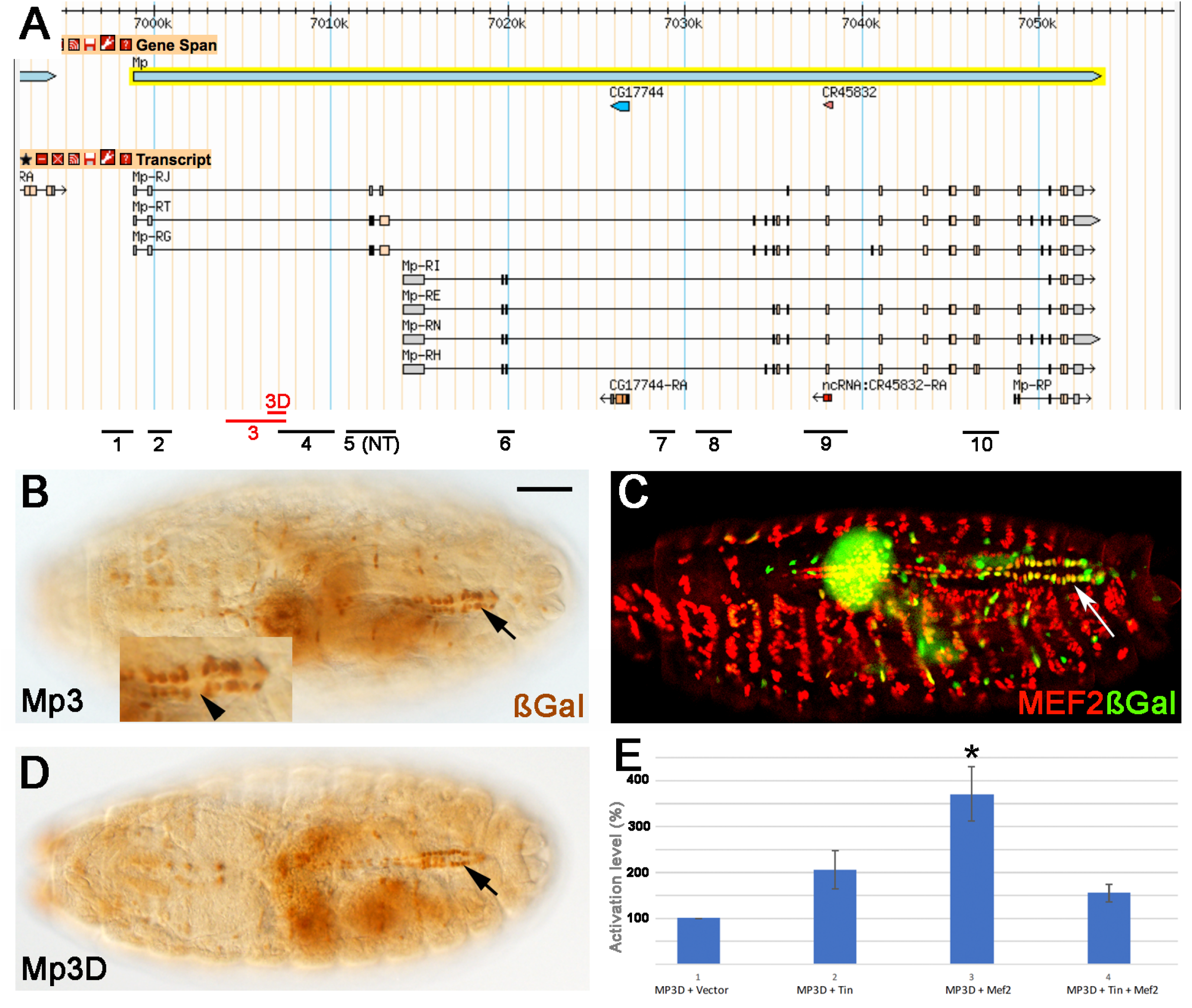
Identification of the Mp enhancer. **A:** GBrowse image from FlyBase.org of the genomic region surrounding *Mp*, including annotation of eight proposed *Mp* transcripts. Black bars at the bottom indicate regions tested in vivo for enhancer activity (NT, not tested). **B:** ßGal accumulation in a stage 16 embryos carrying the *Mp3-lacZ* reporter. Inset shows that not all cardial cells show strong reporter expression in the heart proper (arrowhead). Scale bar, 100µm for B-D. **C:** *Mp3-lacZ* expression overlaps with MEF2 in the heart proper (arrow). **D:** A sub-fragment of Mp3 termed Mp3D also has cardiac enhancer activity in vivo (arrow). **E:** The *Mp3D-lacZ* reporter activated in tissue culture cells by Tin and MEF2.*, p<0.01.

To aid in discovery of the cardiac enhancer for *Mp*, we searched the entire genomic region for the presence of consensus Tin binding sites (5’TYAAGTG; Chen and Schwartz, 1995), and focused upon the areas where two or more sites were located within 300bp of each other. This strategy, although basic, has been successful in identifying cardiac enhancers for both *svp* and *Sur* (Ryan et al 2007; Hendren et al 2007). Using this approach, we identified ten genomic regions to test for enhancer activity (Figure 2A). Each region was amplified by PCR and cloned into a *lacZ* reporter vector, to be used to generate transgenic lines carrying the *Mp-lacZ* constructs. We obtained clones, and subsequently transgenic lines, for nine of these constructs and generated homozygous lines for each. Upon expansion of these stocks, we collected embryos and stained them using anti-ßGalactosidase (ßGal) to visualize reporter activity.

Of all the lines analyzed, only one line, carrying construct Mp3, showed heart-specific expression of the *lacZ* reporter (Figure 2B). The reporter expression was somewhat weak, only appearing in late stage 16 embryos. The reporter also showed expression in some other mesodermal cells likely to be alary muscles and skeletal muscles (Figure 2B) and was not always detected in all myocardial cells (Figure 2B, inset). Nevertheless, expression in the heart co-localized with MEF2 (Figure 2C), indicating that we had indeed identified an *Mp* heart enhancer. A smaller sub-construct of Mp3, designated Mp3D, also showed heart-specific expression, although this was weaker than that for Mp3 alone (Figure 2D; also see below).

To determine if this smaller construct was responsive to the MEF2 and Tin transcription factors, we determined if *Mp3D-lacZ* could be activated in tissue culture by co-transfection with plasmids encoding either MEF2 or Tin. Either factor alone was only mildly capable of inducing *Mp3D-lacZ* expression above background levels, and this activation was only significant for MEF2 (Figure 2E). When we co-transfected both expression plasmids alongside *Mp3D-lacZ*, the activation was intermediate between that for MEF2 or Tin alone, indicating that at least in the context of this enhancer-*lacZ*, these factors did not synergize in their activation of the enhancer (data not shown).

### A minimal Mp cardiac enhancer contains binding sites for both MEF2 and Tin

We next sought to further localize the *Mp* cardiac enhancer, both to localize binding sites for MEF2 and Tin, and to determine if removal of peripheral sequences would generate a minimal enhancer with stronger reporter activity. On the contrary, cardiac enhancer activity seemed to be relatively diffuse across the Mp3 region. We generated transgenic lines carrying seven additional overlapping sub-constructs of Mp3, but only one of these, designated Mp3z, showed reliable expression in the heart, and the level of activity was not noticeably greater than that for the parent *Mp3-lacZ* reporter (Figure 3). *Mp3z-lacZ* was also active in additional cells resembling alary muscles and anterior skeletal muscles.

**Figure 3:**
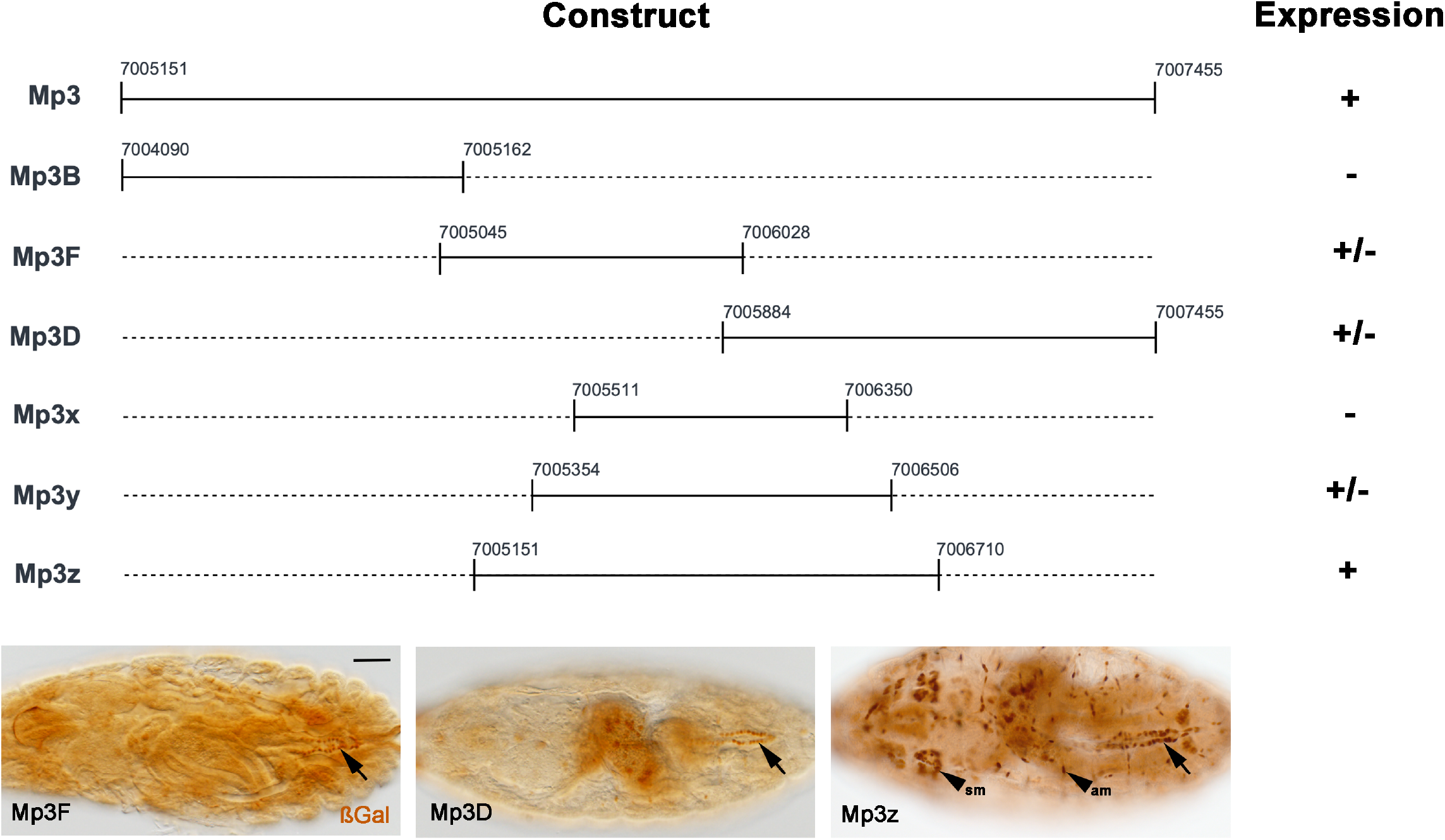
Expression of *Mp3* enhancer-*lacZ* sub-constructs in vivo. Embryos carrying the indicated *Mp-lacZ* constructs were stained for ßGal accumulation in the heart. +, expression was observed at levels comparable to *Mp3-lacZ*; +/-expression was irregular in the heart, with some cells showing expression and others not; -, no cardiac reporter expression observed. Representative images are show at the bottom for Mp3F, Mp3D and Mp3z. Scale bar, 100µm. Genomic coordinates correspond to release 6 of the *Drosophila melanogaster* genome.

The Mp3z DNA is 1560bp in size, within which we searched for consensus binding sequences for both Tin and MEF2 (5’-YTAWWWWTAR; Andres et al 1995). This identified two consensus MEF2 sites (M1 and M2) and three consensus Tin sites (T1-T3). To determine the ability of these sequences to interact specifically with their cognate transcription factors, we performed electrophoretic mobility shift assays (EMSA), where short radioactively-labeled sequences corresponding to each site were combined with MEF2 or Tin protein, and the complexes resolved by native gel electrophoresis. Since M1 and T1 were in close proximity to each other, binding to these sites was tested using a common probe designated M1+T1. The remaining binding sites were tested in isolation.

For MEF2, we used MEF2 protein synthesized in an in vitro transcription/translation reaction, and used unprogrammed lysate as a negative control. For the M1+T1 probe in the presence of MEF2, a complex corresponding to bound probe was observed that was absent from the control lane (compare Figure 4, lanes 1 and 2). The formation of this complex of MEF2 protein with radioactively-labeled probe was competed by the addition of excess wild-type non-radioactive M1+T1 probe (lane 3), but was not competed with the addition of a competitor sequence in which the MEF2 site had been mutated (lane 4). These assays demonstrate that the interaction between MEF2 and the M1+T1 sequence is sequence-specific. A similar set of results was observed for MEF2 interacting with the M2 site (lanes 5-8). We therefore conclude that, at least in vitro, MEF2 can interact strongly with the Mp3z enhancer.

**Figure 4:**
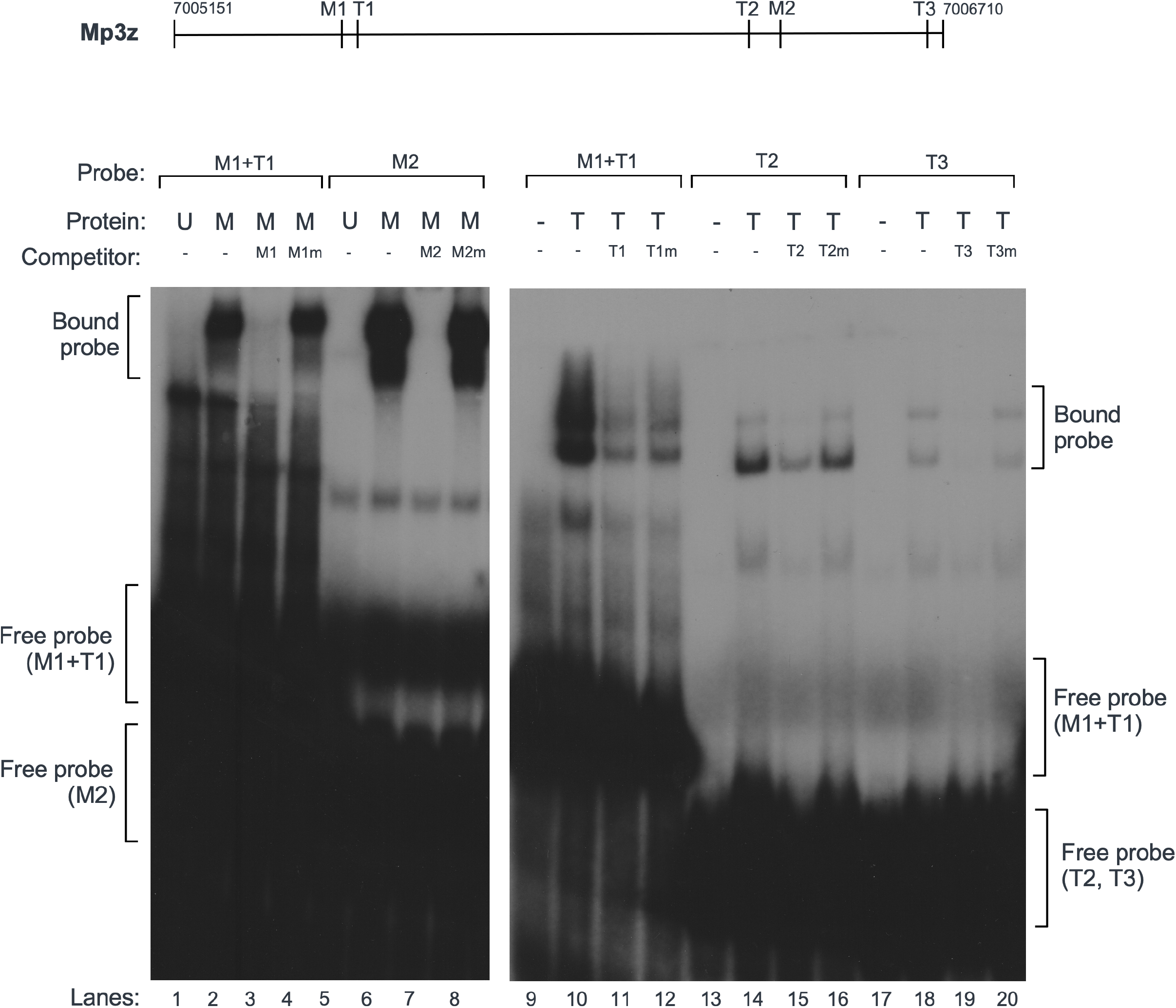
MEF2 and Tin bind to consensus sequences in the Mp3z enhancer. **Top:** schematic of the Mp3z enhancer, with relative locations of MEF2 (M1, M2) and Tin (T1, T2, T3) sites indicated. **Main panels:** electrophoretic mobility shift assays using MEF2 protein (left panel) or Tin protein (right panel). Probe refers to labeled probes used in each set of lanes; Protein refers to either Unprogrammed reticulocyte lysate (U), MEF2 reticulocyte lysate (M), no protein (-), or purified Tin-HD (T). Competitor refers to competitor dsDNAs used in reaction, either wild-type or mutant (lower-case m).

For Tin, we used MBP-Tin protein synthesized in *E. coli*, and for the negative control did not include Tin protein in the reaction (lanes 9, 13 and 17). For M1+T1, we observed a robust interaction between DNA and protein (lane 10), and this interaction was competed with excess unlabeled wild-type M1+T1 (lane 11). However, we also observed a similar, albeit slightly weaker competition in the presence of excess of a version of this probe in which the T1 site had been mutated (lane 12). This suggested that, while Tin could interact with this sequence, a portion of the interaction was non-specific. For the T2 (lanes 13-16) and T3 (lanes 17-20) sites, we observed binding and competition results similar to those observed for MEF2, indicating highly specific interactions between these sequences and Tin.

Overall, DNA binding assays provided strong biochemical support for the hypothesis that MEF2 and Tin each bind directly to the *Mp* gene to activate its transcription.

### Requirement of Tin and MEF2 sites for Mp enhancer activity

In a final experiment, we determined if the integrity of the MEF2 and Tin sites was necessary for Mp3z enhancer activity. We generated transgenic animals carrying *Mp3z-lacZ* constructs in which the MEF2, Tin, or MEF2 plus Tin sites were mutated and assessed reporter expression. Interestingly, mutation of the MEF2 sites resulted in an increase in reporter expression in the posterior of the cardiac tube, and this expression also expanded anteriorly from the heart proper into the aorta (compare Figure 5, panels A and B). This result suggested that the MEF2 sites might overlap additional regulatory sequences whose function is necessary for *Mp* expression specifically in the heart. We address this in greater detail in the Discussion below.

**Figure 5:**
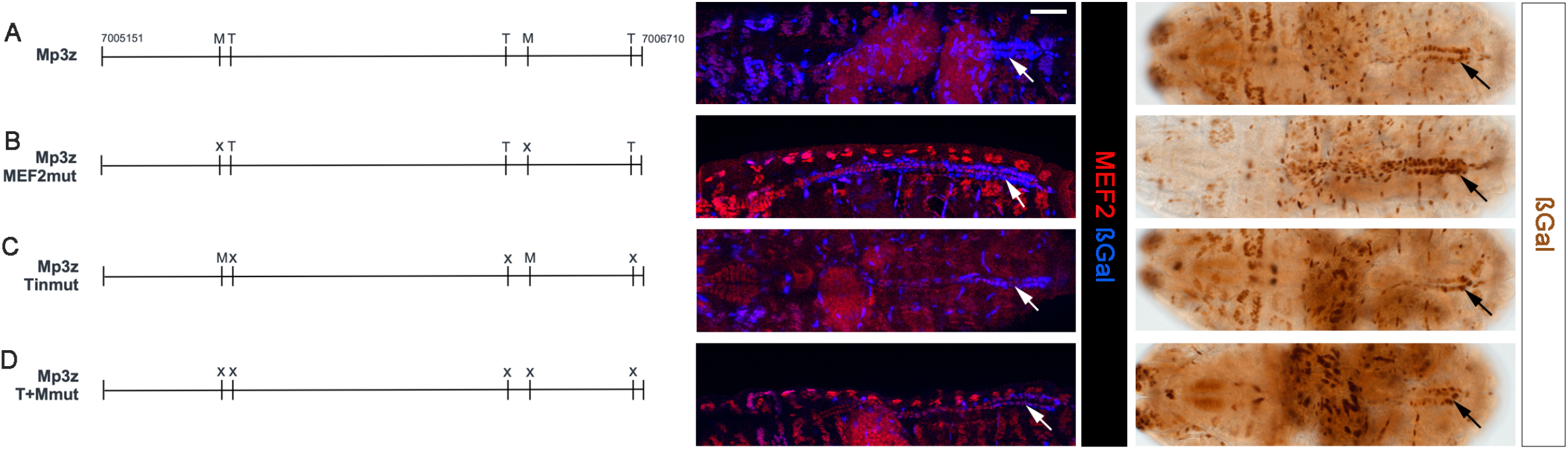
Activity of Mp3z-mutant enhancer-*lacZ* constructs in vivo. The indicated Mp3z constructs at left were fused to a minimal promoter-*lacZ* and tested for enhancer activity in transgenic animals in vivo. Reporter expression was assessed in both fluorescently-stained embryos (center panels) and immunohistochemically (right panels). Note that, as compared to the wild-type reporter-*lacZ* in the heart (**A**, arrow), mutation of the MEF2 sites resulted in increased and expanded ßGal accumulation (**B**). By contrast, mutation of the Tin sites resulted in a strong reduction in reporter expression(**C**). Mutation of both MEF2 and Tin sites also resulted in reduced reporter expression (**D**). Scale bar, 100µm. Genomic coordinates correspond to release 6 of the *Drosophila melanogaster* genome.

Mutation of the three Tin sites resulted in a strong decrease in reporter expression in the heart (Figure 5C), consistent with a requirement for Tin function in *Mp* expression. Similarly, combined mutation of both the

MEF2 and Tin binding sites resulted in an overall reduction in reporter activity, indicating that the requirement for integrity of the Tin sites could overcome the increased expression that we observed when the MEF2 sites were mutated.

Overall, our data demonstrate that Tin and MEF2 each contribute substantially to heart morphogenesis and their role in this process can be at least partially explained at the mechanistic level through a requirement to activate *Mp* expression in the heart.

## Discussion

In this paper we begin to unravel the transcriptional control of heart lumen formation and expansion. Prior research has demonstrated that the formation of a cardiac tube occurs through the actions of a conserved group of structural proteins, however the manner in which this process is orchestrated is still being uncovered. We demonstrate that the muscle transcription factor MEF2 is necessary in Drosophila to form a proper heart lumen, and that expansion of the posterior region of the heart through activation of *Mp* expression is also dependent upon both MEF2 and Tin. These studies importantly begin to connect transcriptional regulators in the heart with the structural genes that directly impact morphogenesis. These connections provide further insight into how mutations affecting cardiac transcription factors can impact multiple mechanisms during cardiogenesis, resulting in pleiotropic human heart abnormalities.

A role for MEF2 in cardiac morphogenesis in Drosophila is reminiscent of the requirement for murine *mef2c* for proper cardiac looping morphogenesis during embryonic development (Lin et al 1997). Moreover, Drosophila MEF2 has been shown to activate alpha-PS2 integrin expression (encoded by *inflated*) in midgut visceral muscle cells to control proper midgut morphogenesis (Ranganayakulu et al 1995). Therefore, a role for MEF2 in regulating ß-PS2 expression (encoded by *myospheroid, mys*) in the heart is perhaps unsurprising. Whether MEF2 directly activated *mys* expression in any context is yet to be determined.

A striking feature of the cardiac tube in *Mef2* mutant animals is the failure of cardiac growth in the posterior of the heart tube, that we show here to arise at least in part due to the loss of *Mp* expression in *Mef2* mutant animals. We note that, since lumen formation is attenuated by the loss of MEF2 function, the failure of heart expansion may also arise, at least in part, due to by the poor initial lumen formation. Nevertheless, we demonstrate clear roles for MEF2, and Tin, in activation of *Mp* expression.

One aspect of *Mp* expression that we have yet to define is how its transcription is restricted to the posterior two segments of the heart, and we note that the *Mp* enhancer is the first such enhancer identified in Drosophila to have region-specific activity along the cardiac tube. Prior work by ourselves and others has shown that the posterior region of the heart is specified by the homeotic gene *abdominal-A* (*abd-A*) (Lovato et al 2002; Lo et al 2002; Ponzielli et al 2002). Therefore, we hypothesize that Abd-A may bind to the *Mp* enhancer to activate *Mp* expression. It is also possible that Ultrabithorax (Ubx) and other homeotic factors, that are expressed in more anterior regions of the heart tube termed the aorta, may function to repress *Mp* expression. We note that high-throughput assays have shown that both MEF2 and Tin can interact physically with Ubx and Abd-A (Baeza et al 2015; Bischof et al 2018), which might be a mechanism for recruitment of homeotic factors to the *Mp* enhancer.

While much of our molecular and genetic evidence points to positive roles for MEF2 and Tin in regulating *Mp* transcription, there are still some issues that remain to be explained. In tissue culture, Tin was unable to significantly activate the Mp enhancer, and mutation of MEF2 binding sites resulted in ectopic enhancer activity for our in vivo reporter rather than a reduction in enhancer activity as we would have predicted. A possible explanation for this result is that the enhancer is also subject to transcriptional repression: there may be proteins present in S2 cells that attenuate enhancer activity, and in vivo the MEF2 sites may overlap with sites for factors that repress *Mp* expression. Such factors could include regulators accumulating in the aorta and not the heart proper, such as Ubx. Nevertheless, this does not fully explain why strong *Mp-lacZ* expression is still observed when MEF2 binding to the enhancer is abrogated. We reason that there must be additional non-consensus sites in the enhancer with which MEF2 can interact to promote *Mp* expression.

Given the dual requirements for Tin and MEF2 in *Mp* expression, and in particular the close proximity of the T1 (Tin) and M1 (MEF2) binding sites, we carried out several assays to determine if these factors might function together. Indeed, we were able to demonstrate a clear physical interaction between the MADS-MEF2 domain of MEF2 and the homeodomain of Tin, using purified proteins and in the absence of DNA. This is consistent with the documented physical interactions of their mammalian counterparts (Vincentz et al 2008; Lei et al 2020). On the other hand, we did not observe synergistic activation of *Mp-lacZ* expression when both factors were present in tissue culture cells. One possible explanation for these observations is that Tin and MEF2 function sequentially in activation of *Mp*, perhaps through a role for Tin in inducing epigenetic changes that facilitate MEF2 accessibility to the enhancer. We conclude that, at least in some scenarios, Tin and MEF2 may collaborate to impact gene expression, however we have yet to effectively demonstrate it in the context of *Mp* transcription.

## Acknowledgements

We thank Jim Shen for assistance with confocal microscopy, Dr. Rolf Bodmer for the *tin* mutant line, Dr. James Skeath for anti-H15 antibody, and Dr. Roger Jacobs for anti-Dystroglycan antibody. This work was supported by R01 GM124498 from the NIH/NIGMS to RMC and Grant MCB-2205405 from the NSF to CAJ. CB is supported by a predoctoral fellowship from the American Heart Association.

